# Temporally coherent perturbation of neural dynamics during retention alters human multi-item working memory

**DOI:** 10.1101/631531

**Authors:** Jiaqi Li, Qiaoli Huang, Qiming Han, Yuanyuan Mi, Huan Luo

## Abstract

Temporarily storing a list of items in working memory (WM), a fundamental ability in cognition, has been posited to rely on the temporal dynamics of multi-item neural representations during retention. Here, we develop a “dynamic perturbation” approach to manipulate the relative memory strength of a list of WM items, by interfering with their neural dynamics during the delay period in a temporally correlated way. Six experiments on human subjects confirm the effectiveness of this WM manipulation method. A computational model combining continuous attractor neural network (CANN) and short-term synaptic plasticity (STP) principles further reproduces all the empirical findings. The model shows that the “dynamic perturbation” modifies the synaptic efficacies of WM items through STP principles, eventually leading to changes in their relative memory strengths. Our results support the causal role of temporal dynamics of neural network in mediating multi-item WM and offer a promising, non-invasive approach to manipulate WM.

## Introduction

Temporarily holding a sequence of items in working memory (WM) for subsequence planning or action is an essential function in many cognitive processes, such as language, movement control, episodic memory, and decision making (Dell et al., 1997; Burgess and Hitch, 1999; Cowan, 2001; Doyon et al., 2003; Giraud and Poeppel, 2012). Psychological evidence and computational models suggest that this process relies on attentional refresh (Awh et al., 1998; Baddeley, 2003; Camos et al., 2018; Oberauer, 2019) or neural reactivations during retention (Lisman and Idiart, 1995; Horn and Opher, 1996; Compte et al., 2000; Raffone and Wolters, 2001; Siegel et al., 2009; Heusser et al., 2016; Lundqvist et al., 2016; Michelmann et al., 2016; Huang et al., 2018; Liu et al., 2019; Schuck and Niv, 2019). Interestingly, recent studies and computational modellings advocates a hidden-state WM network view that does not demand persistent firing during maintenance (Stokes, 2015; Fiebig and Lansner, 2017; Trübutschek et al., 2017; Wolff et al., 2017). Instead, items could be temporarily held in synaptic weights via short-term plasticity (STP) principles (Fusi, 2008; Mongillo et al., 2008; Barak and Tsodyks, 2014; Mi et al., 2017). In fact, the two seemingly contradictory proposals – activated state and hidden state – could be well combined into a unified framework whereby the reactivation and STP mutually facilitate each other and together subserve WM storage (Masse et al., 2018, 2020; Miller et al., 2018). In this WM network, multiple items, by sharing the same connection to an inhibitory pool, compete with each other in time and take turns to elicit brief bouts of reactivation to strengthen their individual synaptic weights (Fino and Yuste, 2011; Mi et al., 2017). Therefore, the dynamic rivalry between item reactivations and the accompanying slowly decaying synaptic weight changes that accumulate over the retention interval would essentially determine the memory strength of each item in the WM network.

Previous studies have examined the neural coding of items and the top-down modulation of WM representations (Sauseng et al., 2009; Rose et al., 2016; Mallett and Lewis-Peacock, 2018; Van Ede et al., 2018), yet the causal evidence for the role of retention-stage neural network dynamics in mediating multi-item WM is largely absent. In fact, unlike optogenetics tools widely used in animal study, there is yet no time-resolved approach to manipulate the dynamics of WM network in the human brain. Here we developed a novel “dynamic perturbation” approach aiming to modify the relative memory strength of a list of items maintained in WM. Specifically, previous studies show that a task-irrelevant color feature will be automatically bound to the to-be-memorized feature (e.g., orientation) in WM when they belong to the same object (Luck et al., 1997; Johnson et al., 2008; Hyun et al., 2009; Huang et al., 2018). Therefore, by manipulating the ongoing temporal relationship between the continuously flickering color probes that are bound to each WM item respectively, we could presumably apply temporally correlated perturbations to these item-specific neural assemblies in the WM network. Consequently, this temporal perturbation would impact the dynamical WM network in certain way and modulate the ongoing competition between items, leading to the changes in their relative STP-based memory strength and finally the subsequent recalling performance.

Six experiments consistently show that the “dynamic perturbation” approach successfully alters the multi-item WM recalling performance on a trial-by-trial basis. We focused on the recency effect (i.e., better memory performance for recently than early presented item), a typical WM behavioral index to characterize the relative memory strength of a sequence of items (Burgess and Hitch, 1999; Gorgoraptis et al., 2011; Baddeley, 2012; Jones and Oberauer, 2013). First, when applying synchronized continuous luminance inputs to the color probes (“Synchronization” manipulation) during retention, the recency effect is significantly disrupted. In contrast, the recency effect keeps intact when luminance inputs are temporally uncorrelated (“Baseline” condition). Second, when the luminance sequences of the color probes are temporally shifted with each other in an order that is either the same or reversed as the stimulus sequence (“Order reversal” manipulation), they lead to distinct and even reversed recency effect. Finally, we established a theoretical continuous attractor neural network (CANN) with STP effects (Brody et al., 2003; Romani and Tsodyks, 2015; Trübutschek et al., 2017; Seeholzer et al., 2019) to mimic the “dynamic perturbation” course. The simulation successfully replicated all our experiment findings, further corroborating the neural mechanism for our manipulation approach. Taken together, our results constitute new causal evidence for the essential role of STP-based neural dynamics in multi-item WM and provides an efficient approach to manipulate WM in humans.

## Results

### “Dynamic perturbation” approach: temporal association of memory-related but task-irrelevant color probes during retention

Subjects performed a WM task in all the experiments. As shown in Figure 1A, participants viewed two serially presented bar stimuli and needed to memorize the orientations of the two bars (“encoding period”). After 5 sec memory retention, participants rotated a horizontal bar to the memorized 1^st^ or 2^nd^ orientation based on the instruction (“recalling period”). Importantly, the two bar stimuli, in addition to having distinct to-be-memorized orientations, also had different colors. During the “maintaining period”, subjects performed a central fixation task, while two discs that had color feature of either the 1^st^ or the 2^nd^ memorized bars (1^st^ memory-related probe, WM1; 2^nd^ memory-related probe, WM2) were displayed at left or right side of the fixation. The color (red, green, or blue) and the spatial locations (left or right) of the two probes (WM1 and WM2) were balanced across trials.

**Figure 1.**
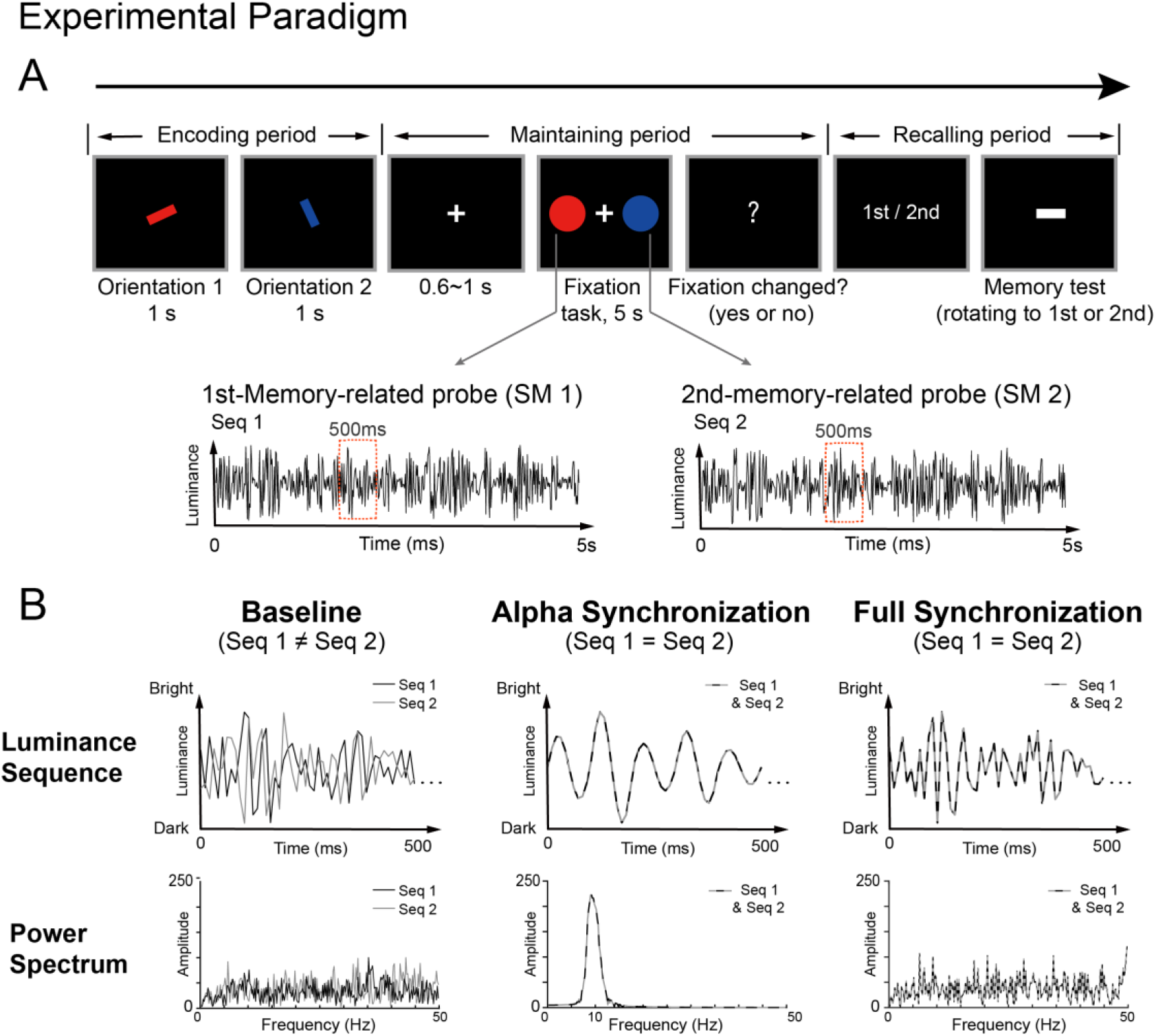
“Dynamic perturbation” approach: temporal association during memory retention. (A) Experimental paradigm. In each trial, in the “encoding period”, two bar stimuli (item 1, item 2) were presented serially at the center. Subjects needed to memorize the orientations of the two bars (1^st^ and 2^nd^ orientation). In the “recalling period”, participants rotated a horizontal bar to the memorized orientation based on the instruction (recalling the 1^st^ or the 2^nd^ orientation). During the 5 s “maintaining period”, subjects performed a central fixation task, while two discs that had color feature of either the 1^st^ or the 2^nd^ memorized bars (1^st^ memory-related probe, WM1; 2^nd^ memory-related probe, WM2) were displayed at left or right side of the fixation. The color (red, blue, or green) and the spatial locations (left or right) of the two probes (WM1 and WM2) were balanced across trials. Throughout the 5 s retention period, the luminance of the two color discs (WM1 and WM2) was continuously modulated according to two 5 sec temporal sequences respectively (Seq1 for WM1, Seq2 for WM2). Note that the luminance sequences were generated anew in each trial. (B) Temporal synchronization of WM1 and WM2 during maintenance. Upper: Illustration of 500 ms segment of the 5 s Seq1 (denoted in dark line) and Seq2 (denoted grey line). Lower: Spectrum of the 5 s Seq1 (dark) and Seq2 (grey). The temporal relationship between Seq1 (black) and Seq2 (grey) varied in one of the three ways (left: Baseline; Middle: A-Sync; Right: F-Sync). In Baseline condition (Left), Seq1 and Seq2 were two independently generated random time series. In A-Sync condition (Middle), Seq1 and Seq2 shared the same alpha-band (8-11 Hz) time series. In F-Sync condition (Right), Seq1 and Seq2 were the same random time series. Note that Seq1 and Seq2 were generated anew in each trial.

Crucially, throughout the 5 sec retention period, the luminance of the two color probes (WM1 and WM2) was continuously modulated according to two 5 sec temporal sequences respectively (i.e., Seq1 for WM1, Seq2 for WM2), the relationship of which varied in certain way. Under Baseline condition, the two luminance sequences (Seq1 and Seq2) were two independently generated random time series, and thus no temporal association between items were introduced (Figure 1B, left). Importantly, we also developed two types of manipulation that aim to apply temporally correlated perturbation – Synchronization (Figure 1B) and Order reversal (Figure 3A) – to the two color probes. For “synchronization” condition, the flickering probes were modulated by the same luminance sequence, either the same random temporal sequence (F-Sync; Figure 1B, right), or the same alpha-band filtered random time series (A-Sync; Figure 1B, middle). Note that including A-Sync condition is motivated by previous findings revealing prominent alpha-band (8-11Hz) activations in WM (Klimesch, 1999; Jensen, 2002). We then examined whether and how the dynamic manipulation impacts the subsequent memory performance, particularly the recency effect, a behavioral index reflecting the relative memory strength of WM items (Burgess and Hitch, 1999; Baddeley, 2012). For Baseline condition when no temporal associations are introduced, we would expect to see intact recency effect, as shown previously (Gorgoraptis et al., 2011; Huang et al., 2018). In contrast, conditions that bring temporally correlated perturbation such as Synchronization or Order reversal would presumably interfere with neural dynamics and impact the recency effect.

It is noteworthy that all the luminance sequences were generated anew in each trial. To quantify the memory performance for the 1^st^ and the 2^nd^ orientation, we used Paul Bays’ model (Bays et al., 2009), a largely used approach that could be implemented through an open-source toolbox, to calculated the target probability (see the precision results in Supplemental Figure 1). We then converted the target probability results using an arcsine square root transformation (Bartlett et al., 1936) to make it normally distributed.

### “Synchronization” manipulation disrupts recency effect (Experiment 1–2)

Twenty-seven subjects participated in Experiment 1 (Figure 2A). First, consistent with our hypothesis, the Baseline condition showed significant recency effect (2^nd^ > 1^st^ item), as typically observed when subjects load a list of items into WM (Gorgoraptis et al., 2011; Huang et al., 2018), suggesting that temporally uncorrelated perturbation would not affect the relative memory strength of WM items. In fact, a control experiment that employed the same paradigm but without flickering color probes presented during retention showed the same recency effect as the Baseline condition (see Supplementary Figure 2). In contrast, when Seq1 and Seq2 are temporally synchronized with each other (A-Sync and F-Sync) by sharing the same luminance sequence, the recency effect was largely disrupted, suggesting their relative memory strengths were successfully modified. Two-way repeated ANOVA (Position * Sync) revealed significant interaction effect (F_(2,52)_ = 3.337, p = 0.043, η_p_^2^ = 0.114), and nonsignificant main effect for either position (F_(1,26)_ = 0.372, p = 0.547, η_p_^2^ = 0.014) or condition (F_(2,52)_ = 0.268, p = 0.766, η_p_^2^ = 0.010), indicating that this dynamic perturbation specifically disrupted recency effect without affecting the overall memory performance. Post-hoc comparisons further confirmed the disruption of recency effect for both A-Sync and F-Sync compared to Baseline (2-way repeated ANOVA; A-Sync and Baseline: interaction effect, F_(1,26)_ = 5.274, p = 0.030, η_p_^2^ = 0.169; F-Sync and Baseline: interaction effect, F_(1,26)_ = 3.503, p = 0.073, η_p_^2^ = 0.119). Moreover, the two synchronization conditions did not show significant difference, indicating their similar disruption effects (2-way repeated ANOVA; A-Sync and F-Sync: interaction effect, F_(1,26)_ = 0.494, p = 0.488, η_p_^2^ = 0.019).

**Figure 2.**
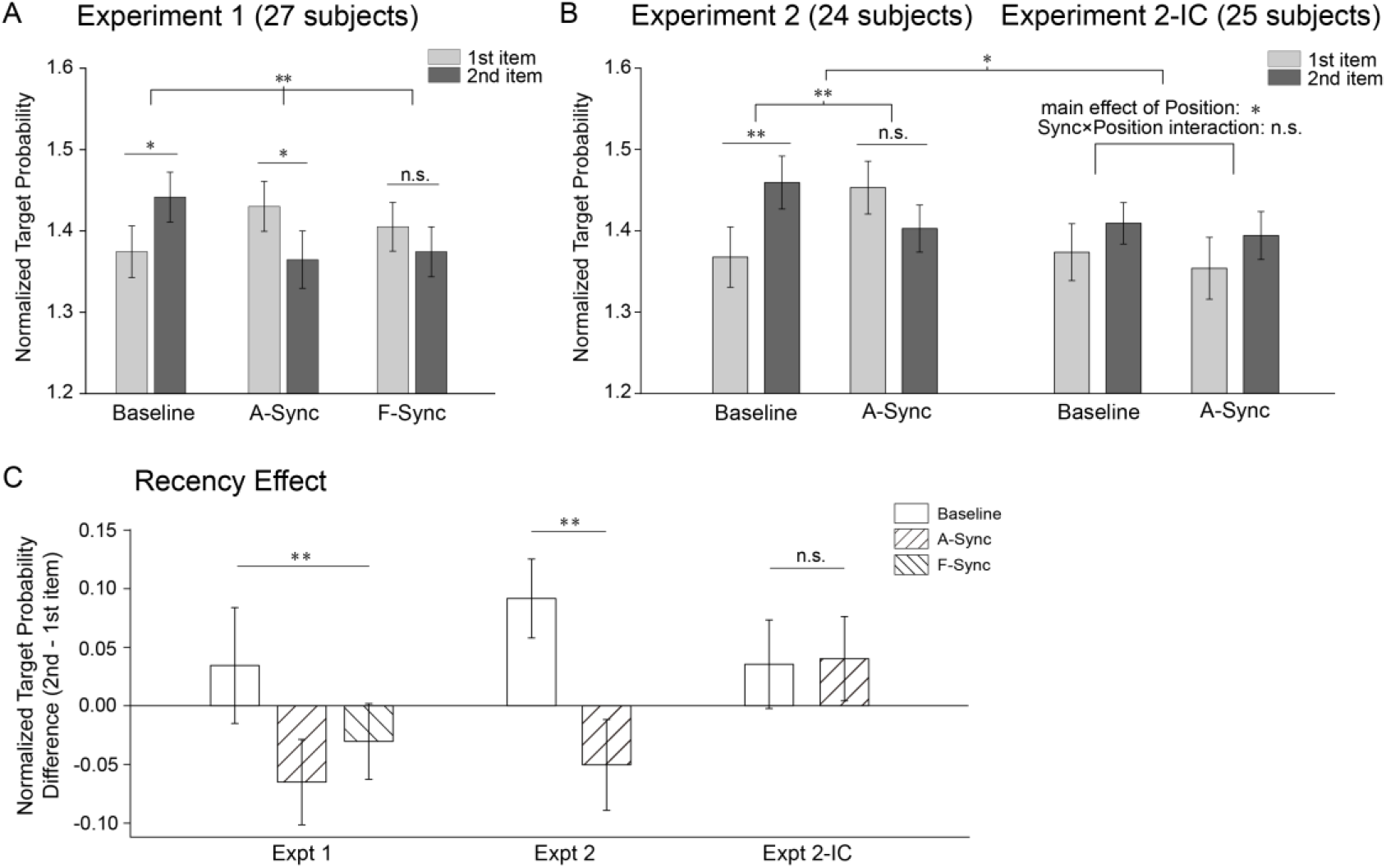
“Temporal synchronization” manipulation disrupts recency effect. A probabilistic mixture model was used to quantify the memory performance for the 1^st^ and the 2^nd^ item. The target probability results were further converted using an arcsine square root transformation to make it normally distributed. (A) Experiment 1 results (N = 27). Grand averaged (mean±SEM) normalized target probability for the 1^st^ (light grey) and 2^nd^ (dark grey) orientation for Baseline, A-Sync, and F-Sync conditions. (B) Experiment 2 (N = 24) and Experiment 2-IC (irrelevant color probes) (N = 25) results. Grand averaged (mean±SEM) normalized target probability for the 1^st^ (light grey) and 2^nd^ (dark grey) orientation for Baseline and A-Sync conditions, in Experiment 2 (left) and Experiment 2-IC (right). (C) Grand averaged (mean±SEM) recency effect (2^nd^ minus 1^st^) for Baseline (white box), A-Sync (rightward checked box), and F-Sync (leftward checked box) conditions in Experiment 1, Experiment 2, and Experiment 2-IC. Noting the significant synchronization-induced recency disruption in Experiment 1 and 2, but not in Experiment 2-IC. (**: p<=0.05, *: p<=0.1).

To further confirm the findings, we ran a separate group of subjects (N = 24) for only the A-Sync and Baseline conditions (Experiment 2). As shown in Figure 2B, the results were well consistent with Experiment 1, such that the Baseline condition showed recency effect (paired t-test, t_(23)_ = −2.721, p = 0.012, Cohen’s d = −0.555), whereas the A-Sync condition did not (paired t-test, t_(23)_ = 1.300, p = 0.207, Cohen’s d = 0.265). Two-way repeated ANOVA (Positions * Sync) revealed significant interaction effect (F_(1,23)_ = 7.493, p = 0.012, η_p_^2^ = 0.246) but nonsignificant main effect for either position (F_(1,23)_ = _0.6_59, p = 0.425, η_p_^2^ = 0.028) or sync condition (F_(1,23)_ = 0.283, p = 0.600, η_p_^2^ = 0.012), further supporting the synchronization-induced disruption effect.

Moreover, to examine whether the disruption effect was specifically introduced by the memory-related probes, we ran another experiment by using memory-irrelevant color probes (Experiment 2-IC, N = 25). As shown in Figure 2B (right), the Baseline and Synchronization conditions now did not show significant difference in recency effect (two-way repeated ANOVA, interaction effect, F_(1,24)_ = 0.006, p = 0.938, η_p_^2^ < 0.001). Two-sample two-way ANOVA (across Experiment 2 and Experiment 2-IC) showed marginally significant interaction effect (Experiment*Position*Sync, F_(1,47)_ = 3.391, p = 0.072, η_p_^2^ = 0.067), further supporting that the manipulation is effective for memory-related probes but not for the non-memory-related ones.

As summarized in Figure 2C, temporal synchronization of the memory-related probes during retention essentially disturbed the recency effect (see similar but nonsignificant trend when spatial locations were synchronized, Experiment 7, Supplementary Figure 3). It is noteworthy that the color probes were completely task-irrelevant in terms of both memory task (memorizing orientations) and attentional task (central fixation task). Moreover, this disruption effect was not accompanied by any overall memory performance changes (i.e., no main effect), indicating that the manipulation does not interrupt memory retention at a general level but indeed specifically modulates the relative memory strength of WM items.

### “Order reversal” manipulation reverses recency effect (Experiment 3-5)

After establishing the efficacy of the synchronization manipulation, we next examined the “order reversal” manipulation. Specifically, the same experimental paradigm as Experiment 1 was used (see Figure 1A), except that the luminance sequences of the two color discs were modulated in a different way. As shown in Figure 3A, Seq1 was a temporally shifted version of Seq2, either leading (Same, upper panel) or lagging Seq2 (Reversed, lower panel) by 200 ms. The selection of 200 ms was motivated by the time constant used in STP neural model (Mongillo et al., 2008; Mi et al., 2017; Trübutschek et al., 2017) and previous neurophysiological studies revealing a central role of theta-band rhythm in organizing memory replay (Lisman and Idiart, 1995; Gyorgy, 2002; Bahramisharif et al., 2018; Huang et al., 2018; Herweg et al., 2020). Two separate groups of subjects participated in Experiment 3 (N = 24; Same, Reversed, Baseline) and Experiment 4 (N=27; Same, Reversed, F-Sync), respectively.

**Figure 3.**
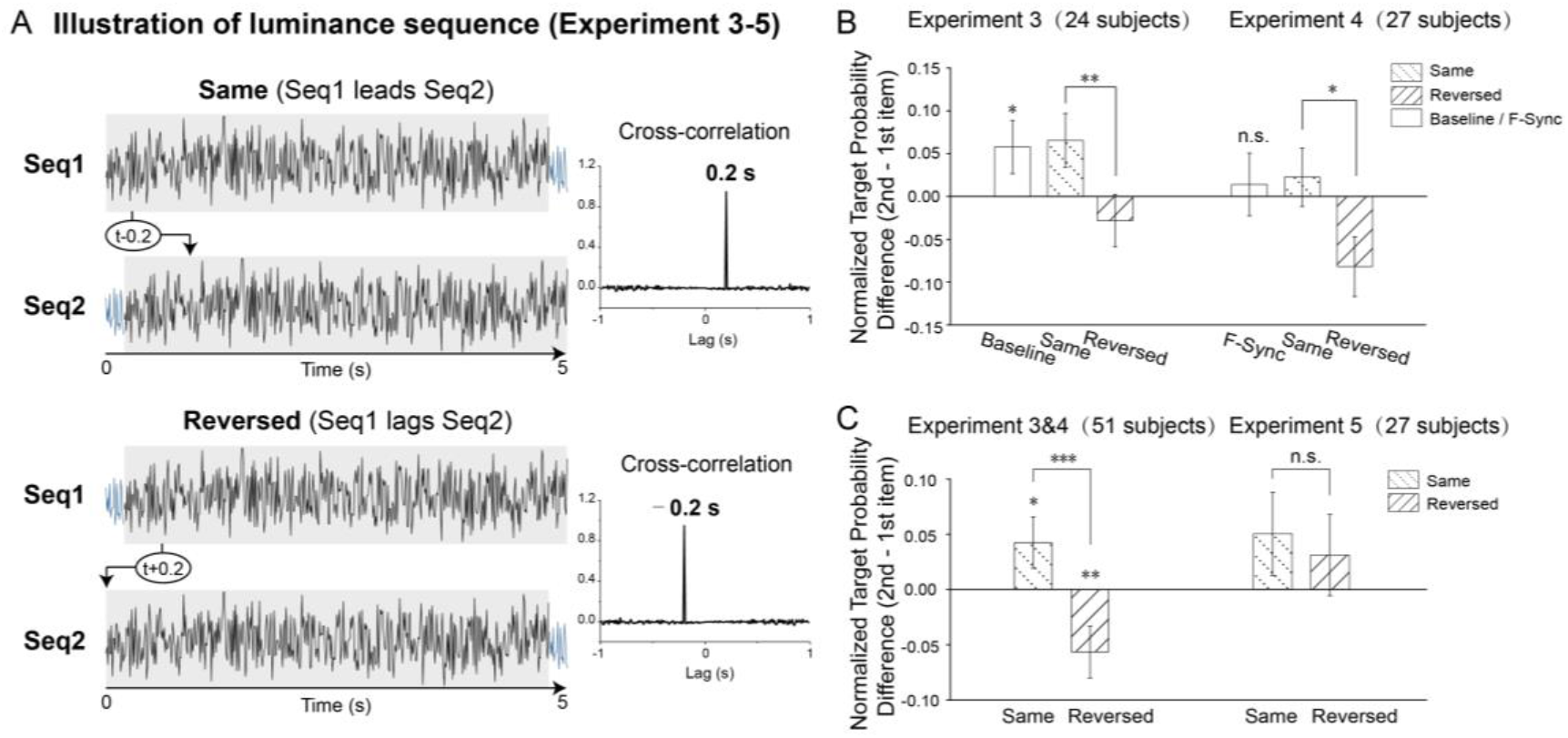
“Order reversal” manipulation disrupts recency effect. **(A)** Experiment 3-5 employed the same experimental paradigm as Experiment 1 (see Figure 1A), except that the luminance sequences (Seq1 and Seq2) were modulated differently. Specifically, Seq1 was a temporally shifted version of Seq2, either leading (“Same-order”, upper panel) or lagging Seq2 (“Reversed-order”, lower panel) by 200 ms. Right: the Seq1-Seq2 cross-correlation coefficients as a function of temporal lag (−1 to 1 s). Note that the luminance sequences were generated anew in each trial. (B) Two separate subject groups participated in Experiment 3 (N=24; Baseline, Same-order, Reversed-order) and Experiment 4 (N=27; F-Sync, Same-order, Reversed-order) respectively. Grand averaged (mean±SEM) recency effect (2^nd^ minus 1^st^) for all conditions in Experiment 3 and Experiment 4 (positive and negative valued represents recency and primacy effect, respectively). Noting the significant decrease in recency effect for Reversed than Same conditions in both experiments. A two-sample two-way repeated ANOVA was performed across Experiment 3 and 4. (C) Left: pooled recency effects (N=51; mean±SEM) for Same-order and Reversed-order conditions across Experiment 3 and 4. Right: Grand average recency effects (N= 27; mean±SEM) for Same-order and Reversed-order conditions in Experiment 5 (500 ms temporal shift). (***: p<=0.01, **: p<=0.05, *: p<=0.1.)

First, as shown in Figure 3B, consistent with previous results (Experiment 1-2), recency effect persisted for Baseline condition (paired t-test, t_(23)_ = −1.835, p = 0.0790, Cohen’s d = −0.375; left panel) and vanished for Synchronization condition (paired t-test, t_(26)_ = −0.366, p = 0.718, Cohen’s d = −0.0704; right panel). Most importantly, the recency effect significantly decreased for Reversed compared to Same condition (paired t-test, Expt. 3: t_(23)_ = 2.184, p = 0.039, Cohen’s d = 0.446; Expt. 4: t_(26)_ = 1.854, p = 0.075, Cohen’s d = 0.357) in both experiments (Figure 3B). Two-sample two-way repeated ANOVA (2*2*2, Experiment*Position*Order) revealed significant Position*Order interaction effect (F_(1,49)_ = 7.543, p = 0.008, η_p_^2^ = 0.133), supporting that the “order reversal” manipulation successfully altered the WM behavior. We further combined the Same and Reversed conditions across Experiment 3 and 4 since the two conditions were exactly the same and derived from two independent subject groups. As shown in Figure 3C (left panel), the pooled results (N = 51) showed clear decrease and even reversal in recency effect for the Reversed condition compared to the Same condition (paired t-test, t_(50)_ = 2.787, p = 0.007, Cohen’s d = 0.390). It is noteworthy that the only difference between the Same and Reversed conditions was the relative order of the two luminance sequences (i.e., Seq1 leads Seq2, Seq2 leads Seq1), yet they led to distinct recency effect. This strongly advocates that the dynamic perturbation in terms of different temporal direction interferes with the WM neural network in opposite ways.

Finally, we assessed whether introducing a temporal lag outside of the STP time window would also influence WM behavior. Instead of 200 ms, Experiment 5 (N = 27) employed a 500 ms temporal lag between Seq1 and Seq2. As shown in the right panel of Figure 3C, the Same and Reversed conditions now did not show significant recency difference (paired t-test, t_(26)_ = 0.331, p = 0.743, d = 0.064). Thus, the dynamic perturbation should be applied within the STP-related temporal window (e.g., short-term depression (STD) time constant) to successfully influence WM.

### Meta-analysis

Finally, to increase the detecting sensitivity and examine the overall effects, we performed a meta-analysis (Comprehensive Meta-Analysis Software, CMA) across all 8 experiments, including Expt. 1 - 5 and Expt. 2-IC in main texts as well as Expt. 6 (probe-absent experiment) and Expt. 7 (spatial location experiment) in Supplementary Materials. As shown in Figure 4, the Baseline condition (Experiment 1, 2, 3, 5, 6, 7, 2-IC) showed significant recency effect (p = 3e^−8^), whereas the Synchronization conditions (Experiment 1, 2, 4, 7) did not (p = 0.424). Furthermore, Baseline and Synchronization conditions (Experiment 1, 2, 7) revealed significant difference (p = 1e^−4^). The meta-analysis on precision measurements showed the same results (Supplementary Figure 4).

**Figure 4.**
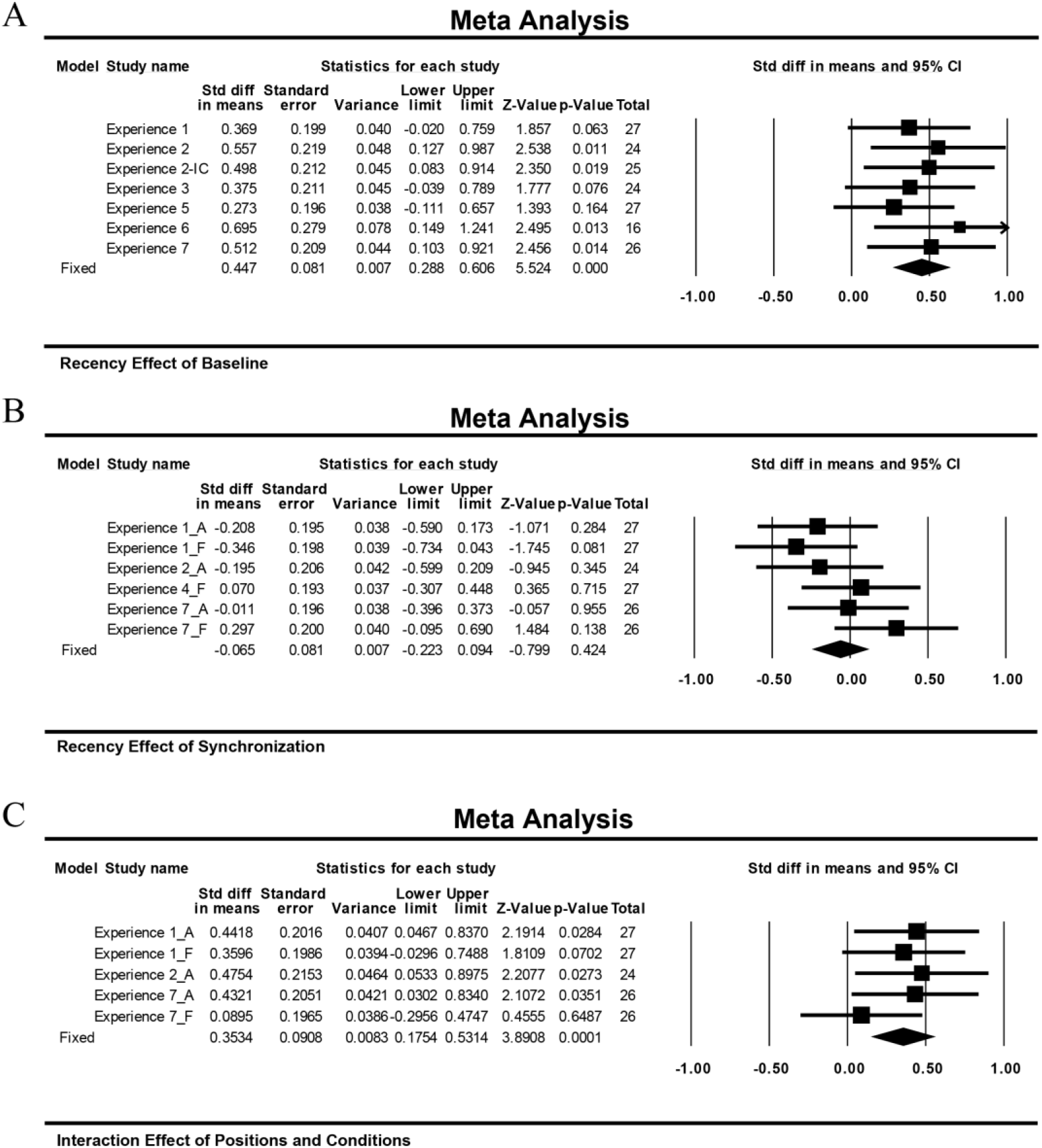
Meta-analysis across eight experiments. Meta-analysis on recency effect across all 8 experiments (Expt. 1-5 and Expt. 2-IC in main text, Expt. 6-7 in Supplementary Mate rials). (A) Significant recency effect for Baseline condition (across Experiment 1, 2, 2-IC, 3, 5, 6, 7). (B) Non-significant recency effect for Synchronization condition (across Experiment 1, 2, 4, 6, 7). (C) Significant difference in recency effect between Baseline and Synchronization conditions (across Experiment 1, 2, 7).

### CANN-STP WM computational model

After confirming the effectiveness of the dynamic perturbation approach on WM manipulation, we further developed a computational model that combines CANN and STP principles to unveil the underlying neural mechanism. CANN has been widely used to model orientation tuning (Ben-Yishai et al., 1995; Blumenfeld et al., 2006) and WM-related network (Brody et al., 2003; Parthasarathy et al., 2019; Seeholzer et al., 2019) in neural systems. STP, referring to a phenomenon that synaptic efficacy changes over time with neuronal activity, is also commonly used to implement WM functions (Mongillo et al., 2008; Barak and Tsodyks, 2014; Mi et al., 2017; Miller et al., 2018). Specifically, as shown in Fig. 5A, in the CANN, excitatory neurons are aligned according to their preferred orientations, connected in a translation-invariant manner. Furthermore, all excitatory neurons are reciprocally connected to a common inhibitory neuron pool (Kim et al., 2017), so that only a single bump-shaped neural response could be generated at any moment. The STP effect is modeled by two dynamical variables, u and x, representing the release probability of neural transmitters (leading to short-term facilitation, STF) and the fraction of resources available after neurotransmitter depletion (leading to STD), respectively (Markram et al., 1998; Mongillo et al., 2008). The instantaneous synaptic efficacy is given by *Jux*, with *J* as the maximum synaptic efficacy. The dynamics of the synaptic efficacy variation is determined by the time constants of *u* and *x*, denoted as *τ*_*f*_ and *τ*_*d*_, respectively. Since STF is dominated in PFC, i.e., *τ*_*f*_ ≫ *τ*_*d*_, we can effectively approximate the STD variable x to be a constant, and only compare the simplified synaptic efficacies *Ju* (see Supplementary Materials and Methods for more details).

**Figure 5.**
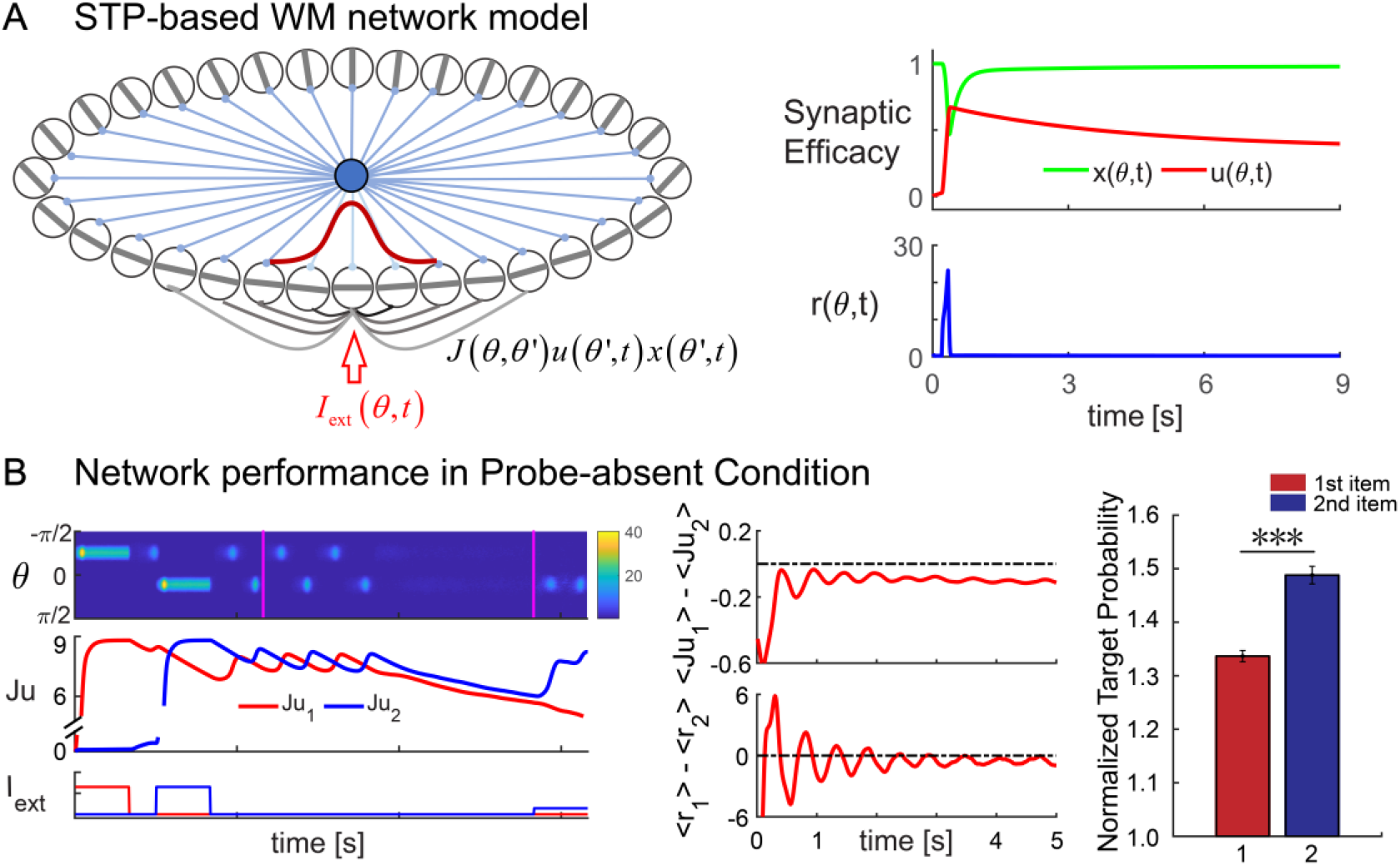
CANN-STP WM network model and simulation of probe-absent condition. **(A) CANN and STP. Left**: Excitatory neurons in the CANN are aligned according to their preferred orientations θ ϵ (−*π*/2, + *π*/2] and they are connected in a translation-invariant manner. All excitatory neurons are reciprocally connected to a common inhibitory neuron pool (blue solid circle). In response to an external input *I*_*ext*_ (*θ*, *t*), the network generates a bump response (red curve) at the corresponding location according to stimulus strength. **Right**: Illustration of STP principle. In response to a transient external input, the neuron encoding orientation at *θ* generates a transient response (firing rate, *r*(*θ*, *t*)), which induces changes in both the neurotransmitter release probability *u*(*θ*, *t*) (red) and the fraction of available neurotransmitter *x*(*θ*, *t*) (green), representing STF and STD effect, respectively. The neuronal synaptic efficacy *Jux*, with *J* as the synaptic weight, will remain at high values for a while, due to the larger time constant for STF than STD (i.e., *τ*_*d*_ = 0.214*s*, *τ*_*f*_ = 4.9*s*). **(B) Simulation results for probe-absent condition. Left**: Individual trial simulation example. Top: Two orientations (1^st^ and 2^nd^) at *θ*_1_ =−*π*/4 and *θ*_2_ =−*π*/4+66° (as examples), respectively, are loaded into the network sequentially and trigger population spikes in the corresponding neuron groups (1^st^ and 2^nd^ neuron group). During retention, the two neuron groups still show response, due to STP, but fall to baseline gradually. In the recalling period, a noisy signal containing partial information of the 2^nd^ memory item (i.e., recalling the 2^nd^ orientation as an example here) is presented and elicits the firing of the 2^nd^ neuron group. Middle: the temporal course of synaptic efficacies of the 1^st^ (red) and 2^nd^ (blue) neuron group throughout the trial. Bottom: the temporal course of external inputs to the 1^st^ (red) and 2^nd^ (blue) neuron group throughout the trial. **Middle:**Grand averaged (mean±SEM, across 20 simulation runs with each having 200 simulation trials) synaptic efficacy difference ⟨*Ju*_1_⟩ − ⟨*Ju*_2_⟩ (top) and firing rate difference ⟨*r*_1_⟩ − ⟨*r*_2_⟩ (bottom) between the 1^st^ and 2^nd^ neuron groups over retention. **Right**: Grand averaged (mean±SEM, across 20 simulation runs) recalling performance characterized by the normalize target probability for the 1^st^ (red) and 2^nd^ (blue) orientation. (***: p<=0.01.)

We then used the CANN-STP WM model to mimic the intrinsic dynamics of neural activities and synaptic efficacies during the delay period as well as the memory recalling performance, for each experimental condition (200 trials in each run, 20 runs for each condition), respectively. Taking one condition as example (Figure 5B), two orientation features are first loaded into the CANN sequentially and evoke two strong transient response (i.e., population spikes). Notably, during retention when stimuli are not present anymore, the synaptic efficacies of the two memory-related neuronal populations would not fall to baseline, and instead, they decay slowly and remain at high values as a result of STP principles. Finally, during recall, a noisy signal containing partial information of a to-be-recalled item is presented to the network, and correct retrieval is achieved if the corresponding neuron group is successfully elicited.

The critical manipulation – flickering color probes – are simulated as continuous and weak inputs, which are endowed with associated temporal characteristics as used in each experiment condition, to the corresponding neural groups during retention (Figure 6, left column, *I*_*ext*_; see more details in Supplementary Materials). These input sequences will perturb the neuronal dynamic through the CANN structure and STP principles and in turn modulate the relative size of the synaptic efficacies of neuron groups (Figure 6, middle column), ultimately determining the memory strength of WM items and the recalling performance (Figure 6, right column).

**Figure 6.**
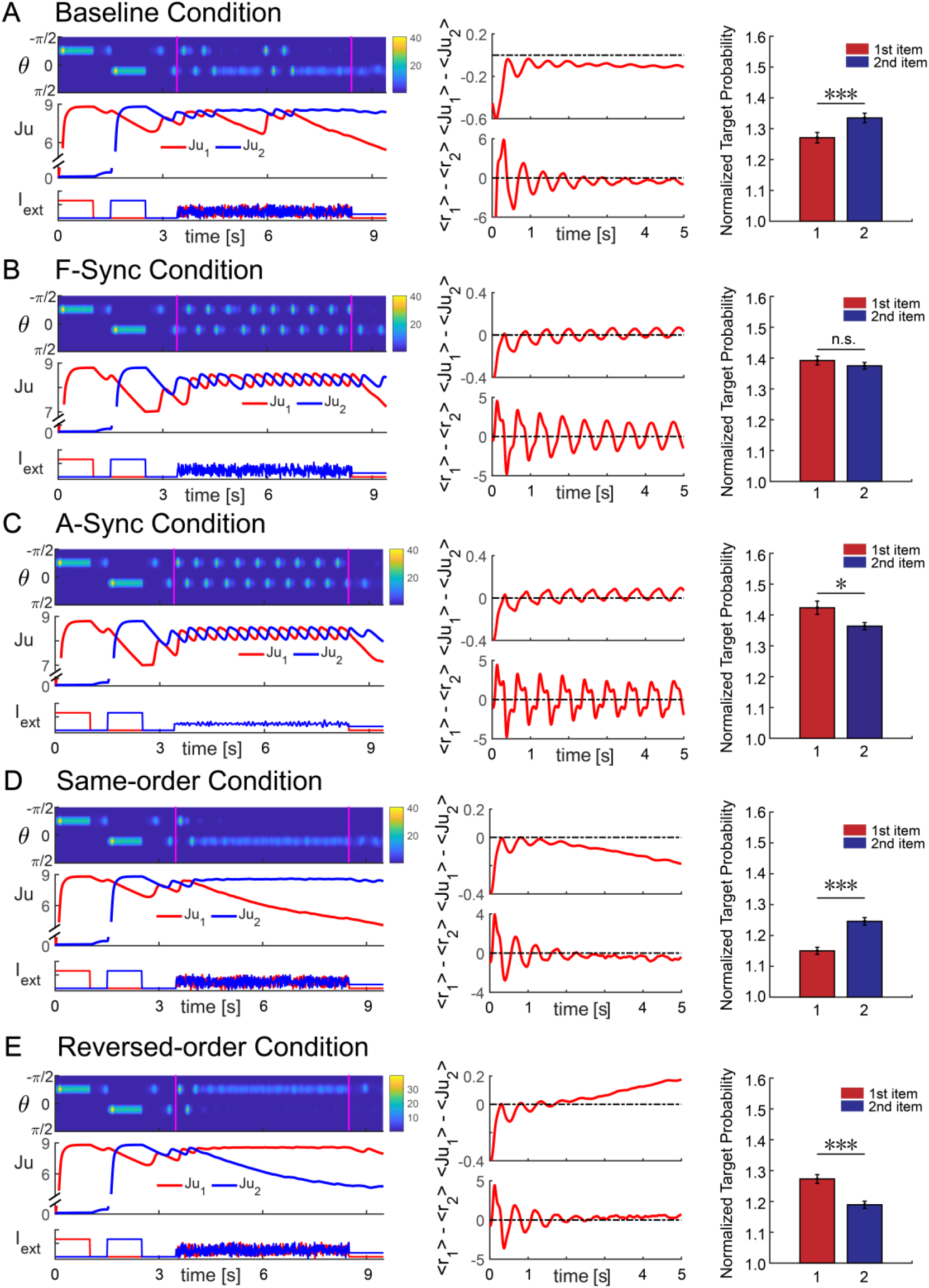
Model simulations for different manipulation conditions. Simulation results for (A) Baseline condition, (B) F-Synchronization condition, (C) A-Synchronization condition, (D) Same-order condition and (E) Reversed-order condition. **Left**: Individual trial example for *θ*_1_ =−*π*/4 and *θ*_2_ =−*π*/4+66°. **Middle**: Grand averaged (mean±SEM, across 20 simulation runs with each having 200 simulation trials) synaptic efficacy difference ⟨*Ju*_1_⟩ − ⟨*Ju*_2_⟩ (top) and firing rate difference ⟨*r*_1_⟩ − ⟨*r*_2_⟩ (bottom) between the 1^st^ and 2^nd^ neuron groups over retention. **Right**: Grand averaged (mean±SEM, across 20 simulation runs) recalling performance characterized by the normalize target probability for the 1st (red) and 2nd (blue) orientation. (***: p<=0.01, **: p<=0.05, *: p<=0.1.). Parameters and mathematical details of probe stimulations are given in Supplementary Materials.

### Model simulations

First, we examined whether our model could reproduce the typical recency effect shown in previous studies (Burgess and Hitch, 1999; Gorgoraptis et al., 2011; Baddeley, 2012; Jones and Oberauer, 2013) as well as our own (Probe-absent experiment, Supplementary Figure 2), when there is no probe stimulation during retention. As shown in Fig. 5B, the loaded items evoke PSs, and the network activity decays to silence rapidly afterwards. Meanwhile, during retention when inputs are removed, the synaptic efficacies of the two neuron groups persist at high values as a result of STP (Figure 5B, left column). Since the two WM items are loaded at different moments, the averaged synaptic efficacy of the early presented item (1^st^ WM item) would be generally smaller than that of the recently presented item (2^nd^ WM item) (Figure 5B, middle column). Consequently, given their difference in synaptic efficacy developed during retention, the 2^nd^ WM item would have larger probability to be successfully retrieved than the 1^st^ WM item (Figure 5B, right column), resulting in the recency effect (paired t-test, t_(19)_ = −7.389, p = 5e^−7^, Cohen’s d = −1.652).

We next examined the Baseline condition, by applying two independent random inputs to the two neuron groups during retention. The probes occasionally trigger PSs in the network (Figure 6A, left column), but they do not essentially disturb the relative size of the synaptic efficacies of the two neuron groups, i.e., ⟨*Ju*_2_⟩ > ⟨*Ju*_1_⟩ still holds, and the mean firing rate for the 2^nd^ item is larger than the 1^st^ item, (⟨*r*_2_⟩ > ⟨*r*_1_⟩) (Figure 6A, middle column). As a result, the Baseline manipulation still leads to recency effect in recalling performance (Figure 6A, right column; paired t-test, t_(19)_ = −3.381, p = 0.003, Cohen’s d = −0.756), consistent with the empirical findings (Figure 2).

For Synchronization conditions, instead of two independent random inputs, the same input series (white noise for F-Sync and alpha-band series for A-Sync) were applied to the two neuron groups (Figure 6BC, left column). Crucially, the synchronized inputs largely disturb the network dynamics and change the relative size of synaptic efficacies, i.e., fluctuating pattern in the synaptic efficacies (Figure 6BC, middle column) as well as the neural activities. Consequently, during recalling (Figure 6BC, right column), the recency effect is largely disrupted (F-Sync: paired t-test, t_(19)_ = 0.971, p = 0.344, Cohen’s d = 0.217; A-Sync: paired t-test, t_(19)_ = 1.905, p = 0.072, Cohen’s d = 0.426), also in line with the experimental observations (Figure 2).

For the “Order reversal” simulation, the input to the neural group that encodes the 1^st^ orientation would either lead (same-order condition) or lag (reversed-order condition) the input to the neural group encoding the 2^nd^ orientation (Figure 6DE, left column). Specifically, the same-order manipulation enhances the relative sizes of the synaptic efficacies of two neuron groups, i.e., enlarged ⟨*Ju*_2_⟩ − ⟨*Ju*_1_⟩ difference and ⟨*r*_2_⟩ > ⟨*r*_1_⟩ (Figure 6D, middle column), finally leading to the recency effect (Figure 6D, right column; paired t-test, t_(19)_ = −6.652, p = 2e^−6^, Cohen’s d = −1.487). In contrast, the “Order reversal” manipulation keeps disturbing the relative size of synaptic efficacies and flips the relationship between them, i.e., ⟨*Ju*_1_⟩ > ⟨*Ju*_2_⟩ and ⟨*r*_1_⟩ > ⟨*r*_2_⟩ (Figure 6E, middle column). This process causes the reversal of the recency effect to primary effect during recalling (Figure 6E, right column; paired t-test, t_(19)_ = 4.363, p = 3e^−4^, Cohen’s d = 0.976), again consistent with the experimental results (Figure 3).

Taken together, the CANN-STP WM model successfully reproduces all the empirical results (Figure 5-6). The model demonstrates that the changes on the ratio of synaptic efficacies of neuron groups, which are caused by the dynamic perturbation during the delay period, result in the manipulation of relative memory strength of WM items.

### Model predictions

Our experiments only tested “Order reversal” manipulation using 200 ms and 500 ms time differences (Figure 3). We further used the model to predict how systematic changes in the correlation time window between the two probe sequences, in the same- or reversed-order conditions, would affect the recency effect (Figure 7). First, when the time lag is close to zero, both the same- and reversed-order conditions show disrupted recency effect and tend to shift to a nonsignificant trend of primary effect, similar to “Synchronization” conditions. Second, when the time lag is too large, both conditions show the recency effect, similar to “Baseline” condition. Most interestingly, when the time lag is within [110 ms, 210 ms], a range close to the temporal constant of STD, i.e., (0.5 ~ 1)\**τ*_*d*_, the recalling performance for the same- or reversed-order conditions begin to display divergence, i.e., recency and primacy effects for the same-order and reversed-order conditions, respectively. Therefore, the temporal window for efficient WM manipulation is determined by the STD time constant.

**Figure 7.**
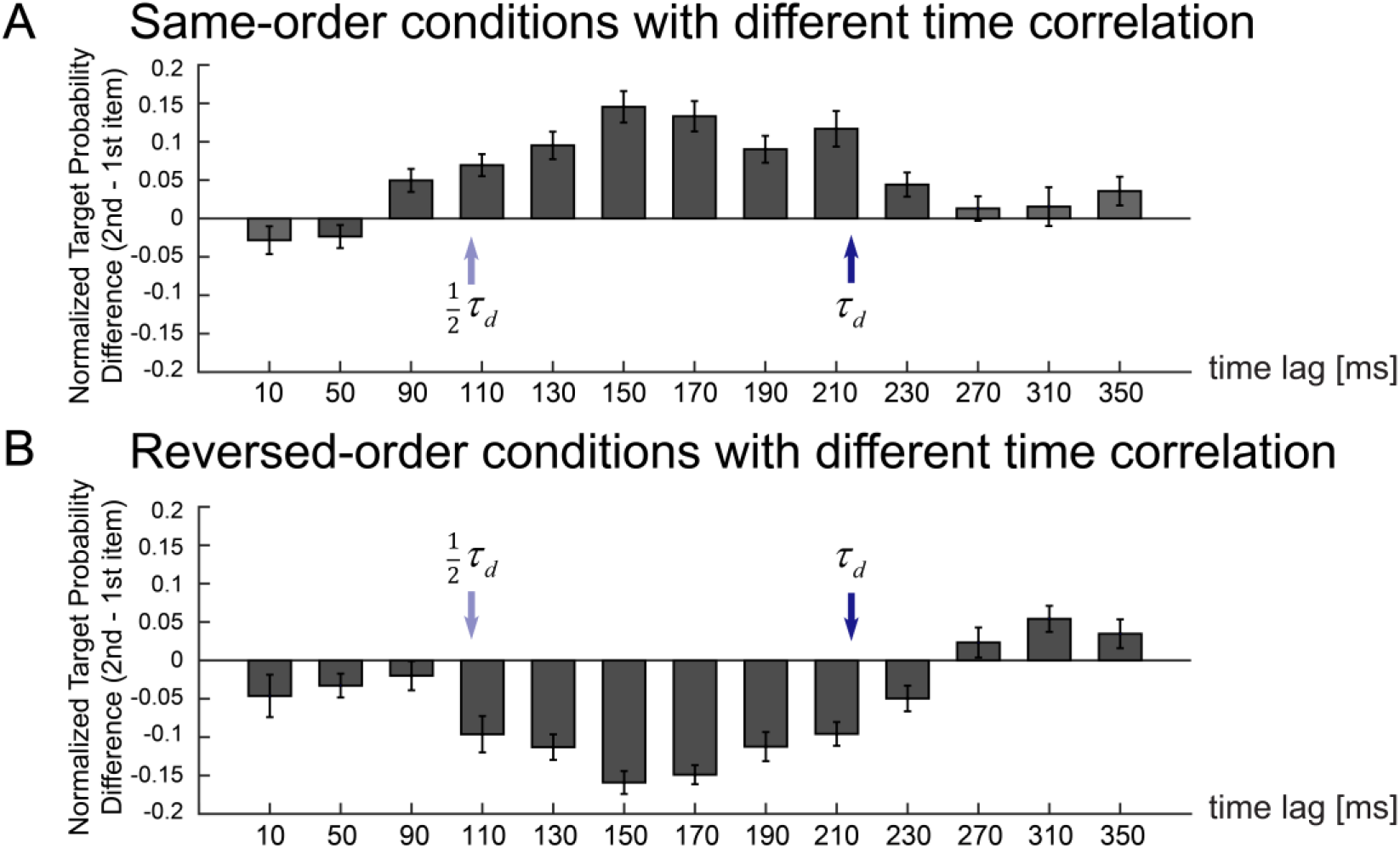
Model predictions. Grand averaged (mean±SEM, across 20 simulation runs with each having 200 simulation trials) normalized target probability difference between the 1^st^ and 2^nd^ orientation (2^nd^ minus 1^st^) at varying time lags (10-350 ms) for (A) same-order and (B) reversed-order conditions. τ_*d*_ (STD time constant) and 0.5τ_*d*_ are marked by dark blue arrows (214 ms) and light blue arrows (107 ms), respectively. Note that when the time lag is close to zero (similar to Synchronization), both the same- or reversed-order conditions show disruption in recency effect, whereas when the time lag is large (similar to Baseline), both conditions show recency effect. Only when the correlation time is in the range of [110 ms, 210 ms], the two conditions show divergence, i.e., recency effect for same-order condition and primacy effect for reversed-order condition. Parameters are given in Supplementary Materials.

## Discussion

We developed a “dynamic perturbation” approach to manipulate the relative memory strength of a list of items held in human WM, by interfering with their neural dynamics in a temporally correlated or uncorrelated way. Six experiments confirm the effectiveness of this method in multi-item WM manipulation. A computational model incorporating CANN and STP principles further reproduces the empirical findings. Importantly, the model demonstrates that this “dynamic perturbation” induces changes in the relative synaptic efficacies of WM items and eventually causes the modulation of their relative memory strength. Taken together, the temporal dynamics of WM neural representation, determined by both active-state and STP-based hidden-state, plays a fundamental role in mediating the storage of multiple items. Our results also provide a promising non-invasive behavioral approach to manipulate human WM.

It has been long posited that temporary maintenance of information relies on a refreshing process (Awh et al., 1998; Baddeley, 2003; Camos et al., 2018; Oberauer, 2019). Animal recordings and human neuroimaging studies show memory-related neural reactivations during retention (Skaggs and Mcnaughton, 1996; Compte, 2000; Wang, 2001; Foster and Wilson, 2006; Siegel et al., 2009; Michelmann et al., 2016; Huang et al., 2018; Liu et al., 2019; Schuck and Niv, 2019), supporting an active-state WM view. Meanwhile, neural response are also known to induce changes in the synaptic efficacy via STP mechanisms (Mongillo et al., 2008; Barak and Tsodyks, 2014; Mi et al., 2017; Trübutschek et al., 2017). Specifically, the synaptic efficacy, given its slow temporal properties, could remain for a while to preserve memory information without relying on sustained neural firing, namely a hidden-state WM view (Stokes, 2015; Wolff et al., 2017; Miller et al., 2018). In fact, the two theories could be well integrated into a framework where information is maintained in the synaptic weights between reactivations through STP principles, and the dynamic interplay between the active-state and hidden-state over the delay period ultimately leads to memory maintenance. Interestingly, sustained firing and hidden-state have recently been posited to be involved in manipulation and passive storage of WM information, respectively (Trübutschek et al., 2017; Masse et al., 2018, 2020). Hence, manipulation of stored information might require reinstatement of the hidden-state representation into activated state (Stokes, 2015; Trübutschek et al., 2019; Masse et al., 2020). Here, by applying certain temporal perturbation patterns to interfere with the active-state of the WM network, we modulate the hidden-state via STP principles and alter the memory strength.

Holding a list of items in WM elicits an item-by-item sequential reactivation pattern (Siegel et al., 2009; Heusser et al., 2016; Michelmann et al., 2016; Bahramisharif et al., 2018; Huang et al., 2018; Liu et al., 2019; Schuck and Niv, 2019), which arises from an ongoing competition between items through lateral inhibition and shared inhibitory inputs (Fino and Yuste, 2011; Kim et al., 2017; Mi et al., 2017). During this rivalry process, each item takes turns to reactivate and accordingly strengthen its respective synaptic efficacy which slowly builds up and decays over the retention interval. Therefore, the dynamic competition between items and the resulted memory trace held in the synaptic weights would eventually determine their relative memory strength. Here, the two WM items are perturbed in a temporally correlated manner so that their dynamic competition is modulated, initiating changes in their relative memory strength. In contrast, temporally uncorrelated perturbation, i.e., baseline condition, given its lack of stable influence on the competition between WM items, shows no modulation effects. Moreover, it is worth noting that the luminance sequences used in our experiment were generated anew in each trial, and the manipulation thus does not depend on specific sequences but on the temporal associations between them.

We build a computational model that combines CANN and STP to explore the dynamics of the WM neural network– both neural firing rate (active state) and synaptic efficacy (hidden state) – and how the dynamics leads to memory strength changes. In this model, STF and STD, as two forms of STP, play different functions, i.e., STF serves to hold memory information and STD controls the time window for effective manipulation. The dynamic sequences brought by the flickering probes continuously perturb the network state and lead to changes in synaptic efficacies and memory performance. When items are independently disturbed in time (Baseline), their competition are not affected and so the recency effect keeps intact, whereas synchronous perturbance would instead change the competition, which in turn modulates their relative synaptic weights and disrupts recency effect (Synchronization). In the case of same-order condition, the perturbation strengthens the relative synaptic efficacies for the two consecutively loaded WM items, resulting in the recency effect. In contrast, under the reversed-order condition, the flipped perturbation pattern indeed reverses the relative synaptic efficacies, i.e.., ⟨*Ju*_2_⟩ > ⟨*Ju*_1_⟩ becomes ⟨*Ju*_2_⟩ < ⟨*Ju*_1_⟩, leading to a shift from recency to primacy effect. Notably, in both cases, the temporal correlation of the two probe sequences needs to fall into the STD temporal range for the manipulation to be effective. How to understand this phenomenon? Previous work (Mi et al., 2017), using STP-based mechanism to explain WM capacity, demonstrates that the time separation between consecutive WM is close to the STD time constant. Therefore, when two temporal sequences are applied to perturb the order of memorized items, their correlation time needs to fall into the STD time constant, so that the changes they induce to synaptic efficacy can effectively disturb the retrieval performances of two memory items. Together, our model sheds light on a challenging WM issue and provides promising ways to efficiently manipulate WM from brain’s perspective.

Previous studies, by using behavioral or non-invasive stimulation approaches in the encoding or delay period, have also successfully modulated the recalling performance (Sauseng et al., 2009; Rose et al., 2016; Clouter et al., 2017; Hanslmayr et al., 2019; Martorell et al., 2019). Here, benefiting from using the time-resolved luminance sequences to disturb the WM network dynamics, our study largely differs from previous interference findings in two aspects. First, previous studies are mainly focused on the modulation of general memory performance while our study aims at particularly manipulating the relative memory strength of WM items (i.e., recency). Second, by using temporally correlated luminance sequences, our approach could precisely modulate the ongoing temporal relationship between items (e.g., synchronization, 200 ms or 500 ms order reversal). Generally speaking, our “synchronization manipulation” condition shares similar motivations as associative memory paradigms, during which pairing two stimuli would lead to changes in their association in memory (Shohamy and Wagner, 2008; Duncan et al., 2012; Gershman et al., 2017). Meanwhile, instead of long-term memory, our manipulation modulates working memory on a trial-by-trial basis. In addition, we adopted a bottom-up approach by modulating the luminance of task-irrelevant color probes, thus largely different from previous interference methods.

Finally, recent studies advocate a close connection between WM and attention, and WM has been proposed to be an internal attentional process that shares similar properties and operation principles as attention (Chun et al., 2011; D’Esposito and Postle, 2015; Oberauer, 2019). For instance, the recency effect could be readily explained in terms of attentional gain, such that the lately presented item enters the focus of attention (FOA) and obtains more attention (Oztekin et al., 2010; Oberauer and Lin, 2017). In general, WM and attention are likely to be mediated via largely overlapping neural networks and similar mechanism, making them hard to be fully disentangled. In fact, the memory-related reactivation might also reflect attentional modulation of neural response that would affect subsequent attentional performance (Soto et al., 2005; Olivers et al., 2006; Mallett and Lewis-Peacock, 2018). Similarly, attention-induced response might leave memory traces to affect following memory behavior (Cowan et al., 2005; Rose et al., 2016; Camos et al., 2018; deBettencourt et al., 2019). Moreover, attention has been shown to sample different objects sequentially and rhythmically, also resembling the dynamic reactivation pattern in WM (Jensen et al., 2014; Jia et al., 2017; Fiebelkorn and Kastner, 2019). Therefore, the “dynamic perturbation” might impact the attentional process as well and thus constitutes a promising way to also manipulate attention, planning and decision making.

## Conflicts of interest

The authors declare no conflicts of interest.

## Acknowledgments

We thank Dr. David Poeppel, Dr. Fang Fang, and Dr. Nai Ding for their helpful comments, as well as anonymous reviews to our previous submission. This work was supported by the National Natural Science Foundation of China (31930052 to H.L., 31771146, 11734004 to Y.Y. Mi), and Beijing Municipal Science & Technology Commission (Z181100001518002), Beijing Nova Program (Z181100006218118, Y. Y. Mi), Guangdong Province with Grant No. 2018B030338001.

## Methods

### Participants

One hundred and ninety-six participants (90 males, age ranging from 18 to 26 years) took part in eight experiments (Expt. 1–5 and Expt. 2-IC in main texts, Expt. 6 (probe-absent experiment), Expt. 7 (spatial location experiment) in Supplementary Materials). One subject in Expt. 1, two in Expt. 2, two in Expt. 2-IC, three in Expt. 3, four in Expt. 4, three in Expt. 5, one in Expt. 6 and five subjects in Expt. 7 were excluded due to their overall low memory performance, or self-reported strategies, or not finishing the whole experiment, resulting in 27, 24, 24, 27, 27, 25, 16, 26 subjects for Expt. 1 to Expt. 5, Expt. 2-IC, and Expt. 6-7 respectively. All the participants were naïve to the purpose of the experiments and had normal or corrected-to-normal vision with no history of neurological disorders, and have provided written informed consent before the experiments. All experiments were carried out in accordance with the Declaration of Helsinki and have been approved by the Research Ethics Committee at Peking University.

### Stimuli and tasks

#### Experiment paradigm

Participants sat in a dark room, 58 cm in front of a CRT monitor (Display++ monitor in Experiment 4) with 100 Hz refresh rate and a resolution of 1024 * 768. Subjects performed a sequence memory task, with their head stabilized on a chin rest. Each trial consisted of three periods: encoding, maintaining, and recalling (Fig. 1A). In the “encoding period”, subjects were serially presented with two bars (Experiment 1, 2, 2-IC, 3, 4, 5, 6, 7) displayed at the center. Each bar stimulus lasted for 1 s with a 0.5 s inter-stimulus interval and had different orientation and color features. Subjects were instructed to memorize the orientations of the bar stimuli. In the “maintaining period”, participants performed a central fixation task by monitoring an abrupt luminance change (within 25% of total trials) and reported if they detected changes after the maintenance period. All trials were included into further analysis. Finally, in the “recalling period”, an instruction cue was presented for 1.5 s to indicate which orientation should be recalled (1^st^ or 2^nd^ orientation), and participants needed to rotate a horizontal white bar by pressing corresponding keys to the memorized orientation as precise as possible, without time limit.

#### Experiment 1–2 (“temporal synchronization” manipulation)

In each trial, after a 0.5 s fixation period, participants performed a two-item sequence memory task. The 1^st^ orientation was chosen randomly between 0° and 180°, and the 2^nd^ orientation was generated based on the 1^st^ orientation with the tiled angles of ±17°, ±24°, ±38°, ±52°, ±66°, ±80° (Liu and Becker, 2013). The color of the 1^st^ and 2^nd^ bar was different (randomly selected from red, green, or blue in each trial). During the maintenance period, after a 0.6-1 s blank interval, two task-irrelevant discs (3° in radius) that had the color feature of either the 1^st^ or 2^nd^ memorized bar, referred as the 1^st^ memory-related probe (WM1) and 2^nd^ memory-related probe (WM2) respectively, were displayed at the left or right side of the fixation (7° visual angle) for 5 secs. The color (red, green, or blue) and the spatial locations (left or right) of the two probes (WM1 and WM2) were carefully balanced across trials. Importantly, throughout the 5 s maintenance period, the luminance of the two color discs (WM1 and WM2) was continuously modulated according to two 5 s temporal sequences, respectively (i.e., Seq1 for WM1, Seq2 for WM2), ranging from dark (0 cd/m^2^) to bright (15 cd/m^2^). The Seq1 and Seq2 were generated anew in each trial.

In Experiment 1, the relationship between Seq1 and Seq2 during the maintenance period varied in three ways: Baseline, alpha-band synchronization (A-Sync), or full-spectrum synchronization (F-Sync). For Baseline condition, Seq1 and Seq2 were two independently generated random time series (Seq1≠Seq2). For F-Sync condition, Seq1 and Seq2 were the same random temporal series (Seq1 = Seq2). For A-Sync condition, Seq1 and Seq2 were the same alpha-band (8-11 Hz) time series (Seq1 = Seq2). Experiment 1 had 216 trials in total (72 trials for each of the three synchronization conditions), divided into three blocks. The three synchronization conditions were randomly mixed within each experimental block.

Experiment 2 only examined the Baseline and A-Sync conditions. Experiment 2 had 144 trials in total (72 trials for each of the two synchronization conditions), divided into three blocks. The two synchronization conditions were randomly mixed within each experimental block.

Experiment 2-IC employed the same design as Experiment 2 (Baseline condition and A-Sync condition), except that the color probes during the delay period had memory-irrelevant colors. Specifically, when the to-be-remembered two bars were blue and red in color, the color probes presented in the delay period would be set to yellow and green, and vice versa.

#### Experiment 3–5 (“order reversal” manipulation)

Experiment 3–5 employed the same paradigm as Experiment 1–2, except that the temporal relationship between Seq1 and Seq2 during the maintenance period were manipulated in a different manner. Specifically, Seq1 was a randomly generated temporal sequence, and Seq2 was a temporal shifted version of Seq1. Specifically, Seq1 either leads or lags Seq2 by 200 ms, corresponding to the “same” and “reversed” conditions, respectively, compared to the stimulus sequence. Furthermore, in order to replicate findings of Experiment 1-2, Baseline and F-Sync conditions were added to Experiment 3 and 4, respectively. Experiment 3 included Same, Reversed, and Baseline conditions, and Experiment 4 included Same, Reversed, and F-Sync conditions. We then assessed the behavioral results using a two-sample (Experiment 3 and 4) two-way (Position*Sync) repeated ANOVA analysis to examine the overall manipulation effect. Each condition had 72 trials, resulting in 216 trials for Experiment 3 and 4, respectively.

Experiment 5 only examined Same and Reversed conditions. Moreover, rather than 200 ms lag in Experiment 3-4, the Seq1-Seq2 temporal lag was set to 500 ms in Experiment 5. Each condition had 72 trials, resulting in 144 trials for Experiment 5.

### Data analysis

A mixture-modelling analysis (Bays et al., 2009) was used to quantify the multi-item WM behavioral performance for each of the item, by quantifying the Von Mises distribution variability of the reported error. The three-parameter mixture model calculates the precision of the reports, as well as the respective proportions of target, distractor and guess responses. In the present study, we used the target probability (see the precision estimation results in Supplementary Materials) that has been widely used to estimate memory performance (Bays et al., 2009; Gorgoraptis et al., 2011; Van Ede et al., 2018). Moreover, the target probability was further normalized by using an arcsine square root transformation (Bartlett et al., 1936). The normalized target probabilities were further used to examine the effect of temporal manipulation on multi-item WM.

The Meta-analysis across experiments were performed by using Comprehensive Meta-Analysis Software (CMA) across all 8 experiments (Expt. 1-5 and Expt. 2-IC in main text, Expt. 6-7 in Supplementary Materials). We calculated the recency effect for Baseline condition across Experiment 1, 2, 3, 5, 6, 7, 2-IC, the recency effect for Synchronization condition across Experiment 1, 2, 4, 6, 7, and the interaction effect between Baseline and Synchronization conditions across Experiment 1, 2, 7.

### Computational modelling

#### Continuous attractor neural network model (CANN)

In this model, excitatory neurons that represent specific orientation (*θ*, (−*π*/2, + *π*/2]) are aligned in the network according to its preferred orientation. They are connected to each other (*J* (*θ*,*θ*′) as the synaptic strength) and are further reciprocally connected to a global inhibitory neural pool (*J*_*EI*_ as the synaptic strength). Specifically, for each excitatory neuron at *θ*, its dynamics of synaptic inputs *h*_*E*_ (*θ*, *t*) is calculated as:

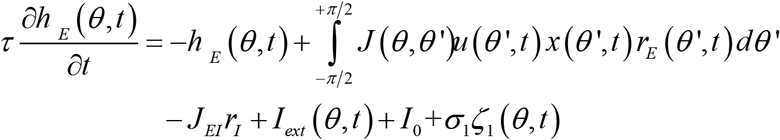

In other words, the dynamics of synaptic current of excitatory neuron at *θ*(*h*_*E*_ (*θ*,*t*)) would be decided by its own dynamics, its stimulus input (*I*_*ext*_(*θ*, *t*)), inputs from other excitatory neurons (*r*_*E*_(*θ*′, *t*)), background activities (*I*_0_ + *σ*_1_*ς*_1_(*θ*, *t*)), and inhibitory inputs (*r*_*I*_). For each neuron, the relationship between firing rate and synaptic current is characterized by *r*(*h*) = *αLn*(1 + exp(*h*/*α*)).

The dynamics of inhibitory neuron is given by 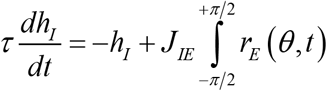, where *r*_*I*_ and *h*_*I*_ denote the firing rate and synaptic current of the inhibitory neural pool, and *r*_*I*_ (*h*_*I*_) = *α* ln(1 + exp (*h*_*I*_/*α*)).

#### Short-term synaptic plasticity (STP)

To model STP effect, *u*(*θ*, *t*) and *x*(*θ*, *t*) are used to quantify the short-term facilitation (STF) and short-term depression (STD) effects, respectively. In fact, *u*(*θ*, *t*) represents the release probability of neural transmitters, and *x*(*θ*, *t*) denotes the fraction of resources available after neurotransmitter depletion, and their dynamics are given by:

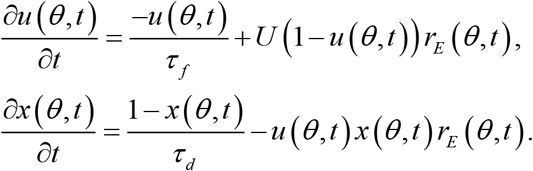

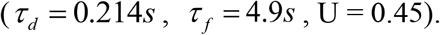

where *τ*_*f*_ and *τ*_*d*_ are the time constants of STF and STD, respectively, and U determines the increment of u due to neuronal firing. We followed the settings in previous STF-dominating WM models, i.e., *τ*_*f*_ << *τ*_*d*_ and a large value of U (*τ*_*d*_ = 0.214*s*, *τ*_*f*_ = 4.9*s*, U = 0.45).

#### Model simulation

We mimicked the whole experimental procedure to do the model simulation. First, during the encoding period, two color bars are loaded into the network sequentially, with each lasting 1 s and the time interval between them being 0.5 s. This is simulated as applying inputs *I*_*ext*_(*θ*_1_, *t*) and *I*_*ext*_(*θ*_2_, *t*) to the CANN, corresponding to the 1^st^ and 2^nd^ orientation features, sequentially. Next, after 0.6-1.4 s, during the delay period, given that the color probes are presumably bound to its associated orientation feature, the 5 sec flickering color probes are simulated by applying a weak, continuous inputs to the CANN respectively. Afterwards, we presented a recalling signal as a loaded item for 1.5 s to retrieve the corresponding memory. Crucially, we mimicked different dynamic perturbation conditions by applying temporal sequences with the corresponding characteristics (e.g., Baseline, Synchronization, Order reversal, etc.) as used in behavioral experiments.

In the recall phase, we used the population vector method to read out the orientation value retrieved by the network, and then followed the same measurement used in psychophysical experiments to calculate the recalling performance. We defined neurons with preferred orientations in the range of |*θ* − *θ*_1_| < 7° as the neuronal group 1, and neurons with preferred orientations in the range of |*θ* − *θ*_2_| < 7° as the neuronal group 2. We also calculated the mean firing rates and the means of synaptic efficacies of two neuronal groups.

The simulation was performed for each experimental condition, respectively. Specifically, 6 pairs of orientation angles with difference (17°, 24°, 38°, 52°, 66°, and 80°) were used for the simulation. For each condition (probe-absent, baseline, synchronization manipulation, same-order, reversed-order, etc.), there were 20 simulation runs and each run contained 200 simulation trials. The results of each run, similar as that of one subject, were then calculated by averaging across the 200 trials that belonged to this run.

For Baseline condition, two independent probe sequences were presented to the 1^st^ and 2^nd^ neuron group during the delay period (Figure 6A). For Synchronization conditions, the same temporal sequences were presented, i.e., seq1 = seq2, during the delay period, either as a white-noise temporal sequence (F-sync condition; Figure 6B) or an alpha-band filtered temporal sequence (A-sync condition; Figure 6C). For the order reversal manipulation, seq1 and seq 2 were a 200 ms temporally shifted version of each other, either seq1 lead seq2 (Same-order condition; Figure 6D) or seq1 lags seq2 (Reversed-order condition; Figure 6E).

For model predictions, the same-order and reversed-order conditions were simulated using varying time lags (10, 50, 90, 110, 130, 150, 170, 190, 210, 230, 270, 310, and 350 ms).

See more model details in Supplementary Materials.

## Notes

### Competing Interest Statement

The authors have declared no competing interest.

### Summary of Updates

Title changed; authors changed; a new computational model added; results reorganized; article largely modified based on the same data.

