## Supplementary Materials for "Temporally coherent perturbation of neural dynamics during retention alters human multi-item working memory"

**Supplementary Figure 1**

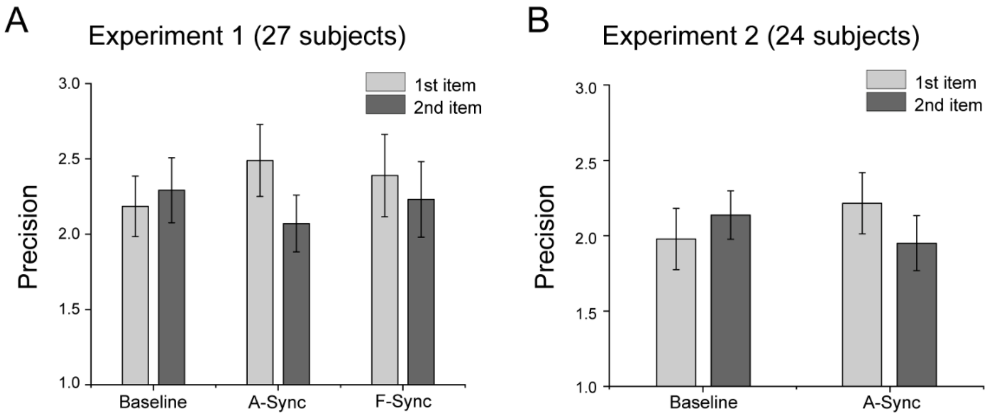


**Precision results (Experiment 1-2)**

Precision results for Experiment 1 and 2, based on the same mixture-modelling analysis. Grand average (mean±SEM) precision results for each item in (A) Experiment 1 (N = 27), Experiment 2 (N = 24), under different "dynamic perturbation" manipulation (Baseline, A-Sync, F-Sync). Although showing similar pattern as the target probability results (Figure 2), the precision results did not reach statistical significance (but see the meta-analysis results, Supplementary Figure 4).

**Supplementary Figure 2**

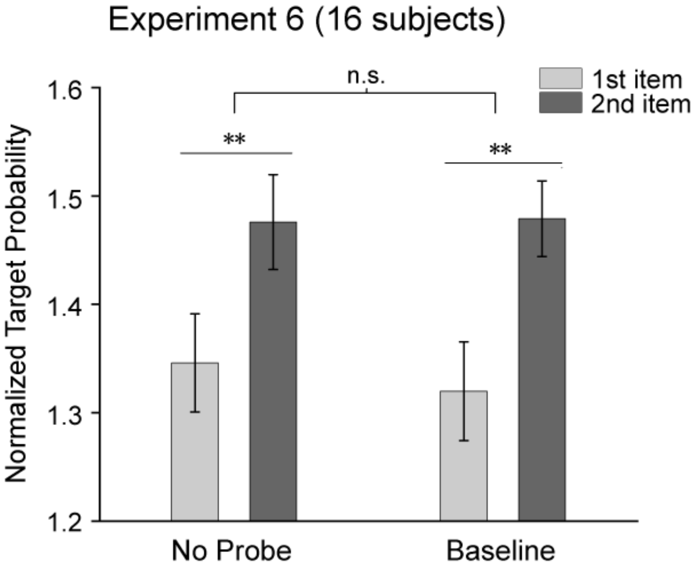


**Probe-absent control experiment (Experiment 6)**

Experiment 6 used the same Experiment 1 paradigm (Figure 1A), except during retention, instead of presenting flickering probes, there was only a central fixation. The probe-absent condition (left) showed similar recency effect (paired t-test, $t_{(15)}$ = -2.426, p = 0.028, Cohen's d = -0.606) as that in Baseline condition (paired t-test, $t_{(15)}$ = -2.793, p = 0.014, Cohen's d = -0.698). Two-way repeated ANOVA revealed significant main effect for position ($F_{(1,15)}$ = 10.789, p = 0.005, $\eta_p^2$ = 0.418), but nonsignificant main effect for condition ($F_{(1,15)}$ = 0.077, p = 0.786, $\eta_p^2$ = 0.005) and nonsignificant interaction effect ($F_{(1,15)}$ = 0.187, p = 0.671, $\eta_p^2$ = 0.012). (**: p<=0.05).

**Supplementary Figure 3**

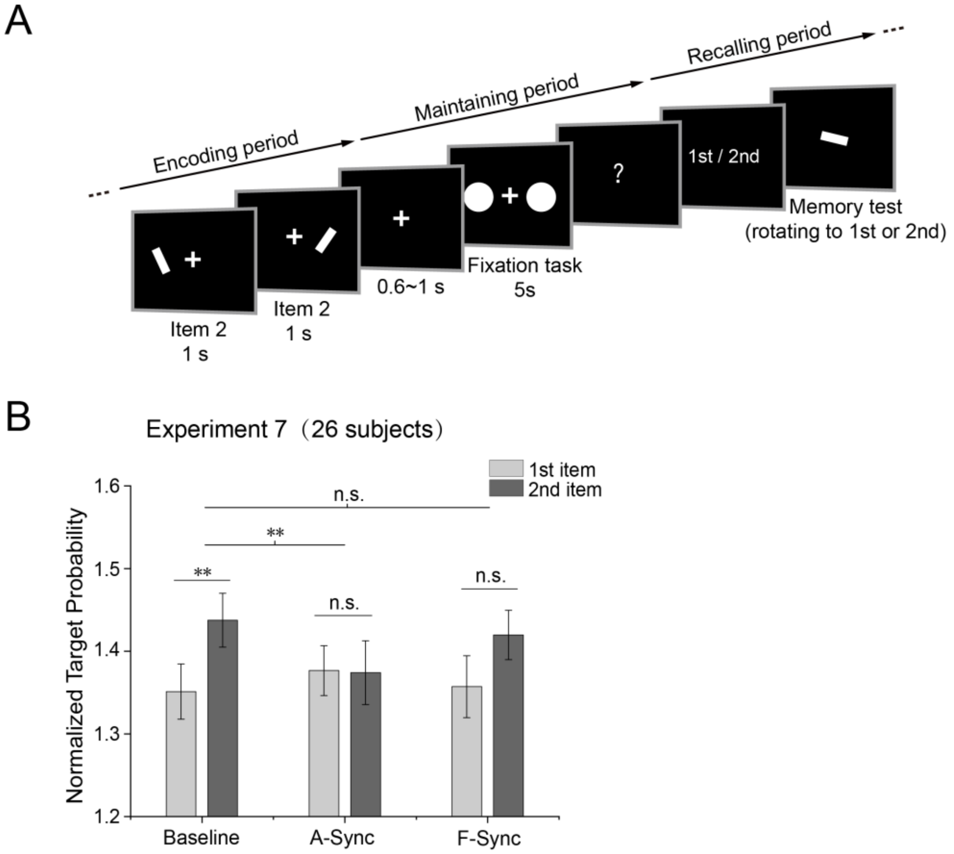


**Temporal synchronization of spatial locations (Experiment 7)**

(A) Twenty-six subjects participated in Experiment 7 (spatial location experiment). In each trial, during the "Encoding period", two white bar stimuli were presented at two different spatial locations serially, and subjects needed to memorize the orientations of two bar stimuli. During the "maintaining period", subjects performed a central fixation task, while at the same time, two white discs were displayed at the spatial locations of either the 1$^{st}$ or the 2$^{nd}$ memorized bars (WM1, WM2). Similar to previous manipulation, the luminance of the two discs were either nonrelated with each other (Baseline) or shared the same time series (A-Sync and F-Sync), and all the luminance sequences were generated anew in each trial. (B) Grand average (mean ± SEM) memory performance for the 1$^{st}$ (grey) and 2$^{nd}$ (black) items in the sequence memory for Baseline, A-Sync, and F-Sync conditions. Temporal synchronization of spatial locations showed a similar trend of synchronization-induced decreased recency effect, but not as strong as manipulation on color probes.

The baseline condition showed typical recency effect (paired t-test, $t_{(25)} = -2.613$, $p = 0.015$, Cohen's $d = -0.513$), whereas the synchronization conditions (A-Sync and F-Sync) did not (paired t-test; A-Sync: $t_{(25)} = 0.057$, $p = 0.955$, Cohen's $d = 0.011$; F-Sync: $t_{(25)} = -1.518$, $p = 0.142$, Cohen's $d = -0.298$). The interaction effect did not reach significance (two-way repeated ANOVA, interaction effect: $F_{(2,50)} = 1.515$, $p = 0.230$, $\eta_p^2 = 0.057$; position main effect: $F_{(1,25)} = 3.787$, $p = 0.063$, $\eta_p^2 = 0.132$; sync condition main effect: $F_{(2,50)} = 0.207$, $p = 0.813$, $\eta_p^2 = 0.008$). (**: $p<=0.05$.)

### Supplementary Figure 4

A

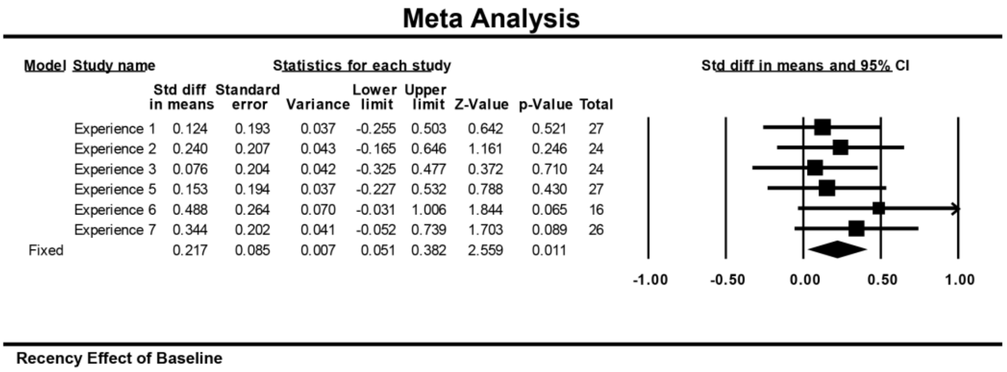


B

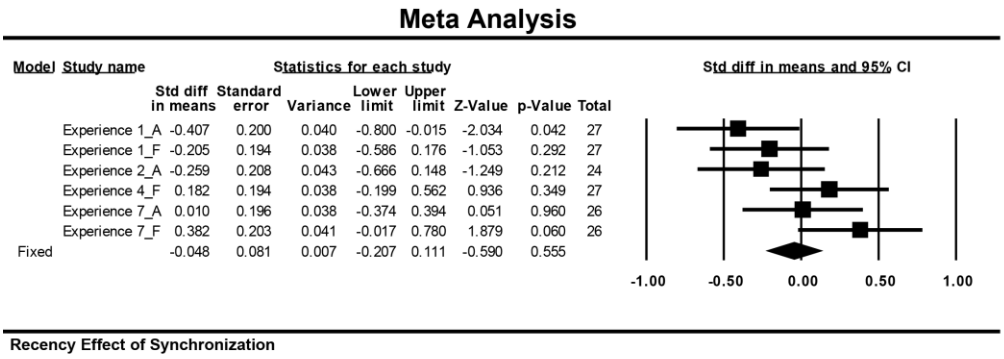


C

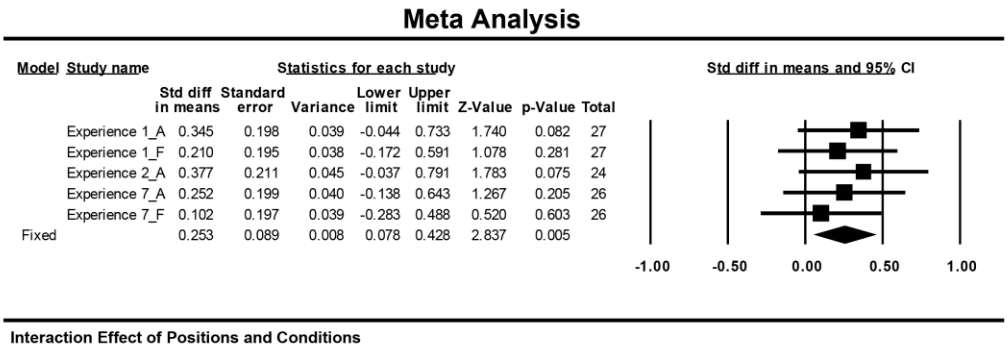


### Meta-analysis on precision results (Experiment 1-7)

Meta-analysis (Comprehensive Meta-Analysis Software, CMA) on recency effect estimated by precision results across all the 8 experiments (Expt. 1-7 and the spatial experiment). Noting the similar synchronization-induced recency disruption effect. (A) Significant recency effect for Baseline condition (across Experiment 1, 2, 3, 5, 6, 7). (B) Non-significant recency effect for Synchronization condition (across Experiment 1, 2, 4, 7). (C) Significant difference in recency effect between Baseline and Synchronization conditions (across Experiment 1, 2, 7).

**Supplementary Information for modeling**

### 1. A continuous attractor neural network with short-term plasticity

We adopt a continuous attractor neural networks model (CANN) with short-term plasticity (STP) to describe how the neural system encodes orientation. Excitatory neurons are aligned in the network according to their preferred orientations, which are in the range of $(-\pi/2,+\pi/2]$ with the period boundary condition. Denote $h_E(\theta,t)$ the synaptic input at time t to neurons at $\theta$, whose dynamics is written as

$$\tau\frac{\partial h_E(\theta,t)}{\partial t}=-h_E(\theta,t)+\int_{-\pi/2}^{+\pi/2}J(\theta,\theta')u(\theta',t)x(\theta',t)r_E(\theta',t)d\theta' -J_{EI}r_I+I_{ext}(\theta,t)+I_0+\sigma_1\zeta_1(\theta,t)$$ Eq.1

where $r_E(\theta,t)$ is the firing rate of excitatory neurons at $\theta$, which is given by $r_E(h_E)=\alpha\ln(1+\exp(h_E/\alpha))$. $I_0+\sigma_1\varsigma_1(\theta,t)$ denotes the background input, with $\varsigma_1(\theta,t)$ the Gaussian white noise of zero mean and unit variance, and $\sigma_1$ is the noise strength. $J(\theta,\theta')$ is the synaptic strength between neurons at $\theta'$ and $\theta$, which is set to be,

$$J(\theta,\theta')=\begin{cases}J\cos[B\times(\theta-\theta')]-J_0, & if\ B\times(\theta-\theta')\in[-ar\cos(-J_0/J),ar\cos(-J_0/J)]\\ -J_0, & else\end{cases}$$

Eq.2

The variables $u(\theta,t)$ and $x(\theta,t)$ quantify the short-term facilitation (STF) and short-term depression (STD) effects, respectively. $u(\theta,t)$ represents the release probability of neural transmitters at time t for neurons at $\theta$, and $x(\theta,t)$ the fraction of resources available after neurotransmitter depletion. The dynamics of $u(\theta,t)$ and $x(\theta,t)$ are given by,

$$\frac{\partial u(\theta,t)}{\partial t}=\frac{-u(\theta,t)}{\tau_f}+U(1-u(\theta,t))r_E(\theta,t),$$
$$\frac{\partial x(\theta,t)}{\partial t}=\frac{1-x(\theta,t)}{\tau_d}-u(\theta,t)x(\theta,t)r_E(\theta,t).$$ Eq.3

where $\tau_f$ and $\tau_d$ are the time constants of STF and STD, respectively, and U determines the increment of u due to neuronal firing. PFC is the main cortical area for working memory (WM), where STP displays the characteristic of STF-dominating, i.e., $\tau_f >> \tau_d$ and a large value of U. This guides us to set the parameters.

All excitatory neurons are reciprocally connected to a global inhibitory neural pool, which induces competitions among neurons, such that only one memory item will be activated at any moment. $r_I$ and $h_I$ denote the firing rate and synaptic current of the inhibitory neural pool, with $r_I(h_I) = \alpha \ln(1+\exp(h_I/\alpha))$. The dynamics of $h_I$ is given by

$$\tau \frac{dh_I}{dt} = -h_I + J_{IE} \int_{-\pi/2}^{+\pi/2} r_E(\theta,t)$$

Eq.4

$I_{ext}(\theta,t)$ is the external input, representing, respectively, the loaded orientation bar signal in the memory loading phase, the flickering probe stimulation in the delay period, and the recalling signal in the memory retrieval phase. Denote $\theta_1$ and $\theta_2$ to be the two memorized orientation values. For the loaded orientation signal and the recalling signal, they are written as,

$$I_{ext}(\theta,t) = \begin{cases} a_{ext}(t)\cos\left(B_{ext}\times(\theta-\theta')\right)+\sigma_2\varsigma_2(\theta,t), & if\ B_{ext}\times(\theta-\theta')\in\left[-ar\cos(0), ar\cos(0)\right] \\ 0 & else \end{cases}$$

Eq.5

where $\theta' = \theta_1$ or $\theta_2$ denote the orientation of a memory item. $a_{ext}(t)$ is the signal strength, $a_{ext}(t) = A_{ext}$ for $t\in[0,T]$, with T is the presentation time of the color bar signal and the recalling signal in the loading and recalling periods, respectively. $B_{ext}$ controls the accuracy of the signal, i.e., the larger $B_{ext}$, the more accurate the signal is. $\varsigma_2(\theta,t)$ is Gaussian white noise of zero mean and unit variance, and $\sigma_2$ the noise strength.

For the probe stimulations, the external input is written as,

$$I_{ext}(\theta,t) = \begin{cases} a^1_{ext}(t)\cos\left(B_{ext}\times(\theta-\theta_1)\right) & if\ B_{ext}\times(\theta-\theta_i)\in\left[-ar\cos(0), ar\cos(0)\right], \\ +a^2_{ext}(t)\cos\left(B_{ext}\times(\theta-\theta_2)\right), & i=1,2 \\ 0 & else \end{cases}$$

Eq.6

where $a^i_{ext}(t)$ is the signal strength, $a^i_{ext}(t)\in[0, A_{ext}]$ for $t\in[0,T]$, with T is the presentation time of the probe signal in the delay period.

For different signals, we choose their accuracies and strengths to satisfy,

$B_{ext}(\text{colorbar}) > B_{ext}(\text{probe}) > B_{ext}(\text{recalling})$, and

$A_{ext}(\text{colorbar}) >> A_{ext}(\text{probe}) >> A_{ext}(\text{recalling})$ .

**2. Parameters used in simulations**

| The CANN Parameters (Eqs.1,2,4) | | | |
|---|---|---|---|
| N | Number of excitatory neurons | 100 | |
| J / $J_0$ | Connection strength between excitatory neurons | 9 | 2 |
| B | Parameter in the connection matrix $J(\theta,\theta')$ | 5.3 | |
| $J_{IE}$ | Connection strength from excitatory neurons to inhibitory pool | 1 | |
| $J_{EI}$ | Connection strength from inhibitory pool to excitatory neurons | 0.28 | |
| $\tau$ | Time constant of synaptic current of excitatory neurons | 20 ms | |
| $\alpha$ | Parameter in the relationship between $r_E$ and $h_E$ | 1.5 | |
| $I_0$ | Background input | -0.9 Hz | |
| $\sigma_1$ | Strength of background input noise in Figure 5B | 2.3 | |
| $\sigma_1$ | Strength of background input noise in Figure 6 and 7 | 0.45 | |
| **The STP Parameters (Eq.3)** | | | |
| $\tau_f$ | Time constant of STF | 4.9 s | |
| $\tau_d$ | Time constant of STD | 0.214 s | |
| U | Increment of release probability | 0.45 | |
| **Parameters of Color-bar Signals (Eq.5)** | | | |
| $A_{ext}$ | Strength of color bar signal | 13 Hz | |
| $B_{ext}$ | Accuracy of input signal | 5.3 | |
| $\sigma_2$ | Strength of noise | 0 | |
| T | Duration of color bar signal | 1 s | |
| **Parameters of Recalling Signals (Eq.5)** | | | |
| $A_{ext}$ | Strength of recalling signal in Figure 5B | 0.95 Hz | |
| $A_{ext}$ | Strength of recalling signal in Figure 6A | 0.8 Hz | |
| $A_{ext}$ | Strength of recalling signal in Figure 6B-E and Figure 7 | 0.48 Hz | |

| | | |
|---|---|---|
| $B_{ext}$ | Accuracy of recalling signal | 2.5 |
| $\sigma_2$ | Strength of noise | 1 |
| T | Duration of recalling signal | 1.5 s |
| **Parameters of Probes Signals (Eq.6)** | | |
| $B_{ext}$ | Accuracy of recalling signals | 4.5 |
| $A_{ext}$ | Maximal probe strength in baseline condition | 3.18 Hz |
| $A_{ext}$ | Maximal probe strength in F-Sync condition | 3.18 Hz |
| $A_{ext}$ | Maximal probe strength in A-Sync condition | 3.18 Hz |
| $A_{ext}$ | Maximal probe strength in same-order condition | 3.2 Hz |
| $A_{ext}$ | Maximal probe strength in reversed-order condition | 3.2 Hz |
| T | Duration of probe stimulation | 5 s |

### 3. The simulation protocol

#### a) The simulation process

The orientations of two color bar signals are set to be the same as in the psychophysical experiments. They are loaded first into the network sequentially, with each of them lasting 1s and the time interval between them 0.5 s. After 0.6~1.4 s, two different flickering color probes with partial information of the memory items (here we consider that the first loaded color bar with orientation $\theta_1$ is binding with red color, and the second color bar with orientation $\theta_2$ with blue color) are presented to the network for 5 s. Afterwards, we present a recalling signal referring to a loaded item for 1.5 s to retrieve the corresponding memory.

#### b) Flickering probe sequences

The sequence strengths of a probe signal in the delay period, i.e., $a_{ext}(t), t \in (0, T)$ are constructed as follows. Mimicking the psychophysical experiments, we first generate a noise sequence X1 of length $T/dt$ (where $T = 5s$ is the duration of delay period, and $\mathrm{dt} = 0.01\mathrm{s}$ is the size of time bin) uniformly distributed in the range of $[0, A_{ext}]$, and calculate its spectrums. We then equalize the spectrum powers of X1 at all frequencies to get a new sequence X2; and finally by scaling the mean of X2 to be $A_{ext}/2$, we get a sequence X3. These sequences and their variations are used to construct the probe strength during the delay period in different conditions.

**1) Baseline condition**

By randomly shuffling the sequence X3, we get another sequence X4. We set X3 to be the sequence of the red-color probe strengths bond with the orientation $\theta_1$, and X4 the sequence of blue-color probe strength bond with the orientation $\theta_2$.

**2) Full-spectrum synchronization condition**

We set X3 to be the sequence of the signal strengths for both the red-color probes (binding with $\theta_1$) and blue-color probes (binding with $\theta_2$) during the delay period.

**3) Alpha-band synchronization condition**

By filtering the sequence X2 with the alpha band, we get a sequence X2'. By scaling the mean of X2' to be $A_{ext}/2$, we get a new sequence X3'. We set X3' to be the sequence of the signal strengths for both the red-color (binding with $\theta_1$) and blue-color (binding with $\theta_2$) probes during the delay period.

**4) Same-order condition**

As shown in Figure 6D of Main Text, we shift the last 200 ms strength sequence of X3 in the front to get X4. We set X3 to be the sequence of the red-color probe strengths bond with the orientation $\theta_1$, and X4 the sequence of the blue-color probe strengths bond with the orientation $\theta_2$.

**5) Reversed-order condition**

The same sequences are constructed as in the same-order condition, but we set X4 to be the sequence of the red-color probe strengths bond with the orientation $\theta_1$, and X3 the sequence of the blue-color probe strengths bond with the orientation $\theta_2$.
